# Cell morphological representations of genes enhance prediction of drug targets

**DOI:** 10.1101/2024.06.08.598076

**Authors:** Niveditha S. Iyer, Daniel J. Michael, S-Y Gordon Chi, John Arevalo, Srinivas Niranj Chandrasekaran, Anne E. Carpenter, Pranav Rajpurkar, Shantanu Singh

## Abstract

Identifying a chemical’s mechanism of action on biological systems is critical for drug development. A compound’s mechanism can sometimes be identified by matching its image-based morphological profile to a well-annotated library. We enhance this approach by incorporating gene representations, demonstrating improved classification performance compared to direct matching methods.

These representations are generated using morphological profiles of cells with artificially altered gene expression. A transformer model is trained to classify gene-compound pairs as true or false interactions, using both gene and compound profiles. The model then ranks likely target genes for unseen compounds.

The strategy performs well for compounds targeting known genes but has limited effectiveness for novel targets, likely due to the current dataset size. However, the performance increase over existing profile-matching methods is notable. Future work with expanded datasets may enhance predictive capabilities, potentially accelerating early-stage drug discovery by improving target identification for novel compounds.

## Introduction

The average cost of introducing a new drug to the market has surpassed a billion USD and continues to rise ^1^. Concurrently, advancements in human genome sequencing have increased the rate of discovery of new diseases and disease subtypes. The current drug discovery approach struggles to match this pace, often focusing on one disease at a time and screening millions of compounds through tailored assays for each disease. Innovations at each stage of the drug discovery workflow are crucial to address this challenge. Modern artificial intelligence and machine learning offer promising solutions, particularly in those stages of drug discovery where large amounts of data are available.

We explore a machine learning strategy to accelerate a crucial step in phenotypic drug discovery: identifying the potential protein targets of a candidate drug. Such compounds may have been discovered to rescue a disease phenotype in cell or organism-based assays, but then require efforts to identify the protein targets they interact with to reveal their mechanism of action. This “target deconvolution” step is often expensive, time-consuming, and inconclusive ^2^.

No single experimental technology can definitively identify a compound’s target ^3^. Proteomics approaches aim to identify unknown protein targets either using pull-down assays or by observing stability shifts in a target in the presence of a compound that binds it. Assay panels, on the other hand, identify engagement of a compound with predefined target classes such as kinases, but can only query a fraction of the druggable proteome.

Recent advancements in high-dimensional biological characterization offer an opportunity to accelerate this process and reduce costs. Image-based profiling assays such as Cell Painting (Fig. 1a) generate information-rich signatures of chemicals by measuring changes in cellular morphology across many parameters, using microscopy images of cells ^4,5^. These assays can profile millions of compounds in screening collections, and all genes in the human genome using genetic perturbations.

**Figure 1.**
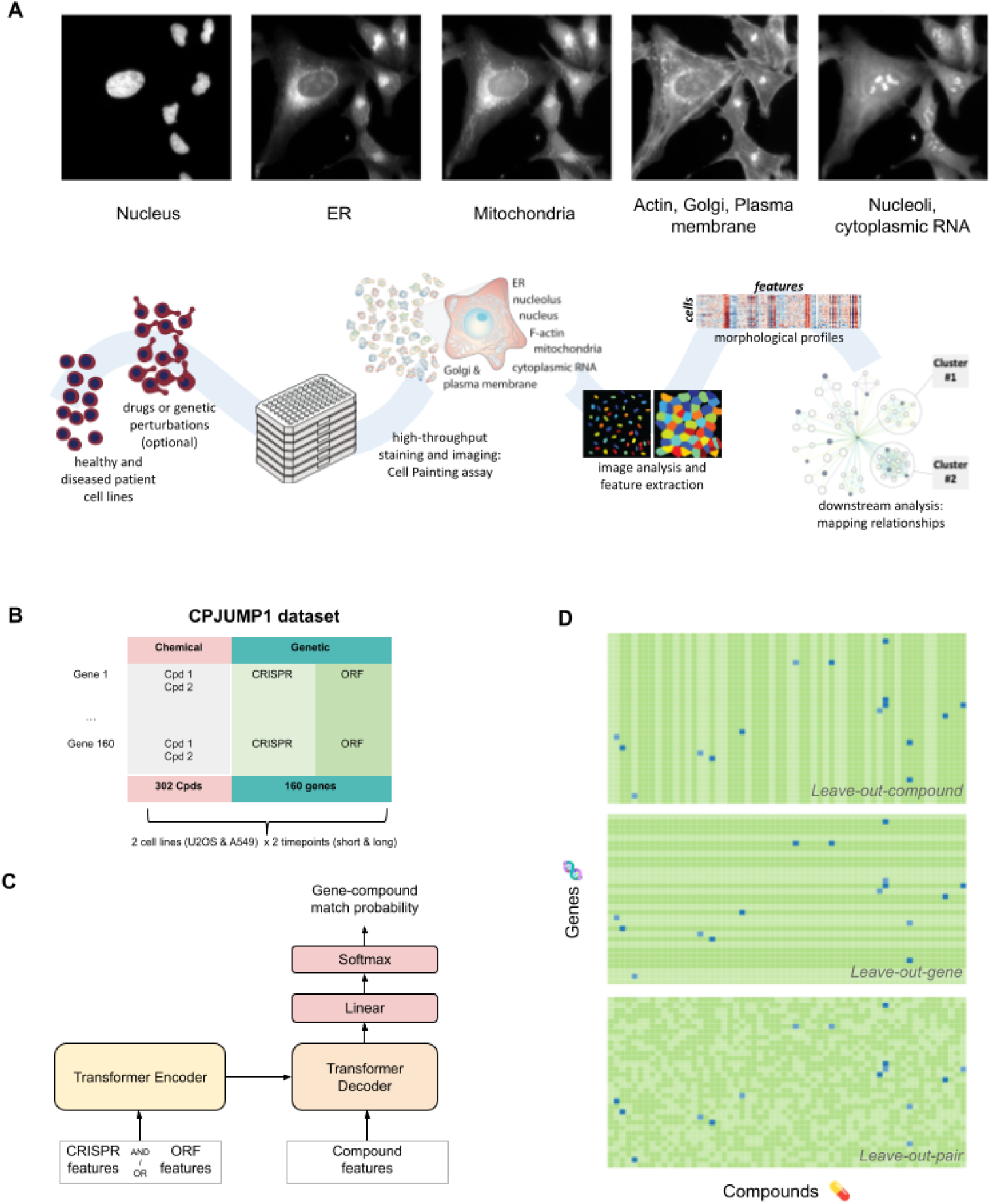
(a) Cell Painting: an image-based profiling assay for generating high-dimensional signatures of chemical and genetic perturbations (adapted from Chandrasekaran et al. ^5^). (b) CPJUMP1 dataset: 302 compounds and 160 genes, each targeted by ≥2 compounds, 2 CRISPR guides, and 1 ORF across 2 cell lines and 2 timepoints. Only a single cell line (U2OS) at a single time point was considered for analysis in this paper; see Methods: Data collection and preprocessing (c) Transformer model: predicts gene-compound match probability from CRISPR, ORF, and compound features. (d) Data split strategies: leave-out-pair, leave-out-compound, and leave-out-gene. Green: no known interaction; blue: known interaction. Darker shades: training set (60%); lighter shades: test (20%) and validation (20%) sets.

Image-based profiling might be used for target deconvolution via two strategies, both based on the hypothesis that perturbing the same targets generate similar phenotypic signatures:

1. Matching query compounds with annotated compounds. Here, profiles of compounds with unknown targets are matched to those of well-annotated compounds.
2. Matching query compounds with perturbed genes. Here, profiles of compounds with unknown targets are matched to those of cells perturbed with a specific genetic reagent, providing evidence that the gene, or other related genes with similar function, encodes the protein target of the compound.

Compound-compound matching has become well-established over the past 20 years, in both the published literature and the pharmaceutical industry ^6^. Compound-gene matching (the only option for compounds with a novel mechanism of action) has only recently been tested across many genes ^7^ and has not yet been widely implemented in practice. Simple similarity matching works well for target classes that manifest strong, broad alterations in cellular morphology ^5^.

However, the signal of target classes with subtle impacts can be drowned out by technical noise. Furthermore, for compound-gene matching specifically, similarity metrics leave much room for improvement because the different perturbation modalities (chemical vs genetic perturbations) are challenging to align ^8^. The main limitation is the dependency on a singular, global similarity metric which fails to capture the subtle variations of different target classes and perturbation modalities. We hypothesized that a learning approach might be superior and improve compound-gene matching.

Here, we use high-dimensional image-derived signatures from Cell Painting to train machine learning models to predict gene-compound “matches”, aiming to overcome this challenge. We present a systematic approach employing a transformer-based model to predict putative compound targets using a benchmarking dataset of 302 compounds and 160 genes.

Despite our modest dataset size compared to typical chemical libraries and the human genome, our initial findings show a significant improvement in target deconvolution, hinting at a promising avenue to reduce screening costs and accelerate drug discovery.

## Results

### Experimental design and methodology for predicting drug targets

This study advances image-based profiling methods for target deconvolution, which can be done by matching profiles of compounds to annotated compounds or, less commonly, perturbed genes ^6,7,9^. We posited that integrating gene morphological profiles into a machine learning framework could enhance the accuracy of predicting compound targets. To explore this, we developed a transformer-based model trained to classify gene-compound pairs as true or false interactions, drawing on the CPJUMP1 dataset ^8^.

### CPJUMP1 Dataset

The CPJUMP1 dataset (Fig. 1b) comprises Cell Painting profiles from chemical and genetic perturbations in two cancer cell lines, U2OS (osteosarcoma) and A549 (lung carcinoma). It includes pairs of compounds and genes likely to form matching relationships based on manual annotation. The chemical perturbations are 302 small molecules from the Drug Repurposing Hub^10^, over a third of which are launched drugs and many others having reached clinical trials. All compounds were profiled at a single concentration of 5μM. Genetic perturbations consist of gene overexpression via open reading frames (ORFs) and gene inhibition through CRISPR-Cas9-mediated knockout, covering 160 genes. See Supporting Information B for further details about the genes and compounds.

Selection criteria for compounds included targeting diverse gene families and ensuring each gene was targeted by at least two compounds while excluding markedly non-selective compounds ^10^. However, it should be strongly noted that we do not expect perfect gene-compound matching in this dataset for several reasons: most compounds are annotated with more than a single target (with ∼21% targeting proteins from multiple gene families), a compound may influence only a single function of a multi-functional protein, the high compound concentration may induce polypharmacological effects not seen at typical pharmacological doses especially for potent clinical compounds like kinase inhibitors, a compound may have an impact only in a certain cell type not used in the study, and the annotations are incomplete and imperfect. Furthermore, the perturbation effects and models are specific to the cancer cell line contexts used.

### Experimental Design

We assessed three baseline approaches for predicting compound targets using this dataset: (a) direct gene-compound profile matching ^7^, (b) compound-compound similarity ^9^, and (c) training a classifier for each gene target ^9^. Our proposed method, by contrast to (b) and (c), incorporates the gene’s morphological profile when assessing whether a compound targets that gene.

Inspired by transformer models from natural language processing, we adopted an encoder-decoder architecture for our biological data. In these models, words are typically represented as dense, low-dimensional vectors called embeddings, which are a specific type of learned representation. In our model, Cell Painting profiles from CRISPR, ORF, and compound perturbations are treated as embeddings. This model takes genetic perturbation embeddings as the source sequence and compound embeddings as the target sequence, processed through the encoder and decoder respectively (see Fig. 1c); thus, the language of genetic perturbations is “translated” to the language of chemical perturbations. The transformer’s output passes through a fully-connected and a softmax layer to generate a binary prediction probability.

We evaluated the model under three strategies of splitting the set of the 302 × 160 gene-compound pairs, in all cases creating training (60%), validation (20%), and test (20%) sets (Fig. 1d):

1. Leave-out-compound: Split the 302 compounds such that the validation and test sets did not contain any compound-gene pairs for *compounds* seen during training.
2. Leave-out-gene: Split the 160 genes such that the validation and test sets did not contain any compound-gene pairs for *genes* seen during training.
3. Leave-out-pair: Randomly split gene-compound pairs.

In all strategies, each gene-compound pair is exclusive to one set. Pairs are labeled *positive* if the gene’s protein product is a known target of the compound, and *unknown* otherwise. The leave-out-compound and leave-out-gene strategies simulate predicting targets for novel compounds or genes, respectively. The dataset was designed to cover diverse gene families to mitigate leakage from highly related genes.

We reported model performance using Precision@R for each split strategy, where R represents the number of true positives ^11^ (see *Methods*). We chose Precision@K as our primary metric because it quantifies the proportion of correct predictions among the top-ranked candidates—a key consideration for target prediction tasks. All reported results are on the held-out test set. By comparing strategies, we aimed to identify the most effective approach for predicting compound targets using morphological profiles.

### Cell morphological representations of genes enhance prediction of drug targets

To assess the model’s ability to predict gene-compound interactions for previously unseen compounds (but where another compound targeting the same gene may have been seen), we tested the *leave-out-compound* strategy. In this scenario, the model is tasked with determining if a gene-compound pair is a true match (i.e., positive), despite the compound being absent from the training data. We evaluated three baseline approaches within this framework.

The first baseline, direct gene-compound profile matching (Figure 2A, “gene-cpd simil.”) yielded very low performance. Although this approach had previously identified some hits ^7^, our results demonstrate the signal is relatively weak compared to other approaches, likely due to differing morphological effects of the two perturbation types that normalization techniques alone may not suffice to align.

**Figure 2.**
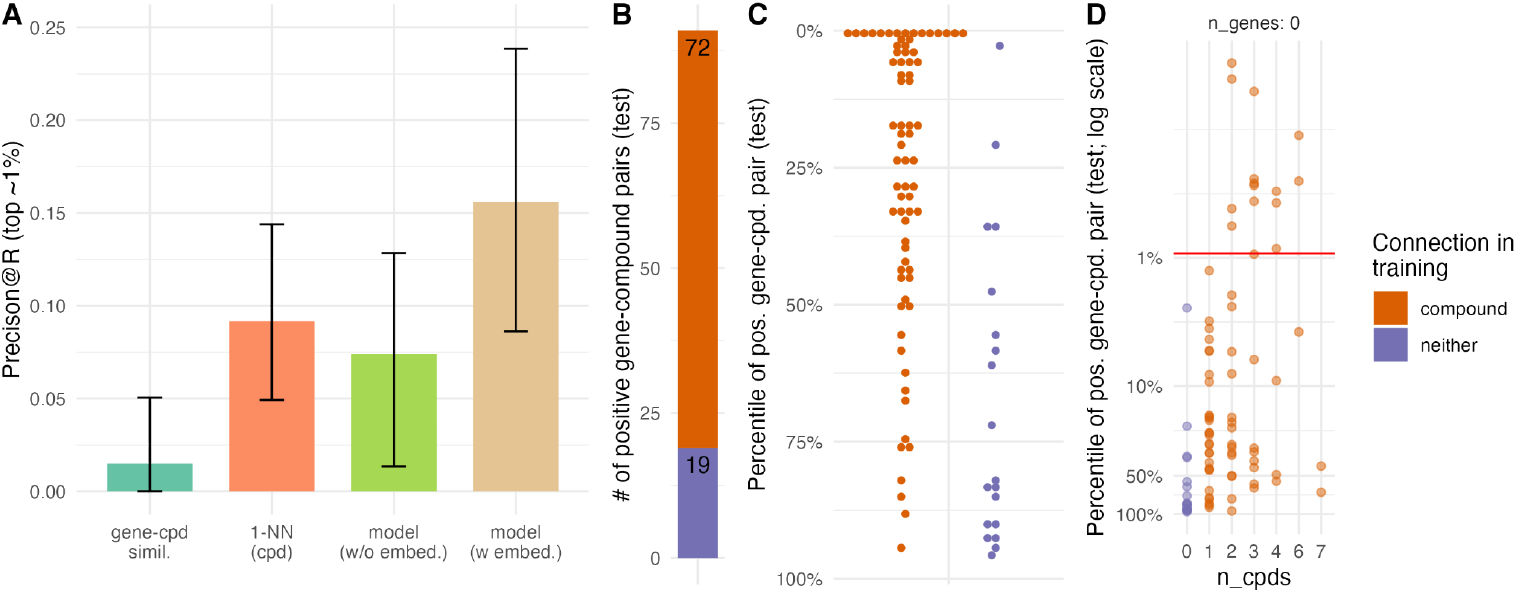
Leave-out-compound performance; retrieving the correct gene target for a “novel” compound (one that has not been seen during training). (A) Proposed method (“model (w. embed.)”) vs. baselines: gene-compound similarity (“gene-cpd simil.”), 1-NN compound classifier trained for each gene (“1-nn cpd”), transformer without embeddings (“model (w/o embed.)”). Error bars: 95% CI. (B) Counts of two categories of positive gene-compound pairings from the test set: positive gene-compound pairings where the test gene is connected to a training compound (“compound”) or not (“neither”). Test compounds are never in the training set. (C) Model predictions of positive gene-compound pairings from test set (as percentiles): correct classification is likelier (top percentile) if the test gene is connected to a training compound (red) vs. not (blue). (D) Model predictions of positive gene-compound pairings from test set (as log-scaled percentiles) by number of training compounds connected to the test gene: performance improves with more connections.

The second baseline, 1-NN compound-compound similarity ^9^ (Figure 2A, “1-NN (cpd)”), labels a test gene-compound pair as positive if the compound’s nearest neighbor in the training set interacts with the gene. This outperformed direct matching but cannot predict interactions for genes unseen during training, limiting its utility for novel targets.

Our proposed model (Figure 2A, “model (w embed.)”), which incorporates genetic perturbation profiles, surpassed both baselines. However, we wondered whether the high performance of the model could be explained by the structure of the transformer framework alone, and if it might be using the gene embeddings simply as gene identifiers or labels rather than using them for the morphological information they contain. To test this, we evaluated a third baseline where we shuffled the embeddings, effectively reducing them to gene labels, and used the transformer model for prediction (Figure 2A, “model (w/o embed.)”). By doing so, we essentially trained a separate classifier for each gene within the transformer model. Notably, the transformer using shuffled embeddings (i.e., gene labels) performed no better than the simpler 1-nearest neighbor (1-NN) model based on compound similarity only. This observation suggests that the morphological information captured by the gene embeddings, rather than their mere use as gene identifiers, is crucial for enhancing target prediction performance.

Examining the model’s performance in detail, we first note that for 72 out of 91 positive gene-compound pairings in the test set, the test gene was connected to a training compound (red; Fig. 2b), i.e. the test gene was seen in training in a positive pair. The top-scoring positive gene-compound pairings are enriched for seen genes; for unseen genes, predictions are essentially random (Fig. 2c). This suggests that the model can predict target genes for unseen compounds but not the subset of those that target novel genes. We also examined the impact of the number of compounds targeting a gene during training; as expected, having more compounds targeting a gene improves performance for other compounds targeting that gene (Fig. 2d).

Overall, our model outperforms baselines in the leave-out-compound strategy, demonstrating the importance of gene embeddings in predicting the target of unseen compounds. However, it has limitations in predicting targets for compounds that target novel genes, a topic we explore next.

### Predicting compound targets for novel genes is diminished

The leave-out-gene strategy, in which the model is tasked with identifying targets for compounds that target unseen genes, reveals a significant reduction in the model’s predictive performance (Fig. 3). This stark contrast to the leave-out-compound strategy’s results prompted an investigation into the underlying factors contributing to this asymmetry.

**Figure 3.**
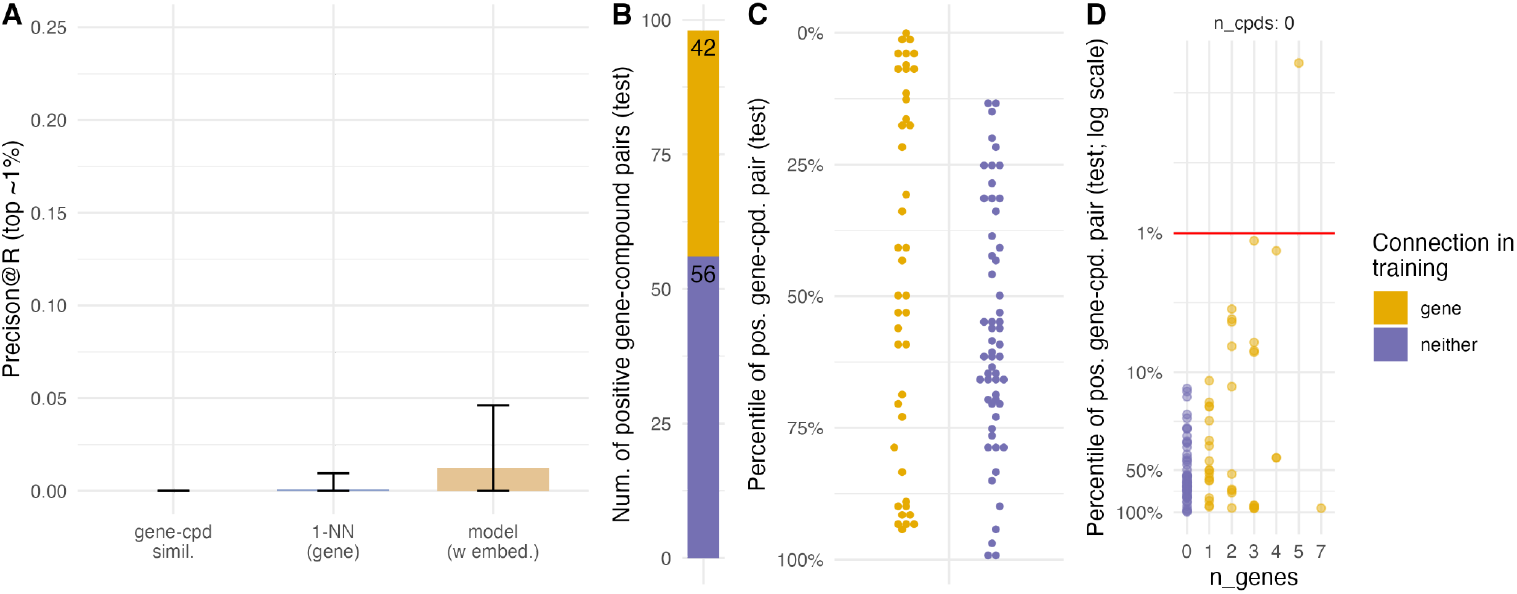
Leave-out-gene performance; retrieving the correct target for compounds having a “novel” gene target (one that has not been seen during training). (A) Proposed method (“model (w. embed.)”) vs. baselines: gene-compound similarity (“gene-cpd simil.”) and 1-NN gene classifier trained for each compound (“1-NN gene”). Error bars: 95% CI. (B) Counts of two categories of positive gene-compound pairings from the test set: gene-compound pairings where the test compound is connected to a training gene (“gene”) or not (“neither”). Test genes are never in the training set. (C) Model predictions of positive gene-compound pairings from test set (as percentiles): correct classification is generally poor and is no likelier if the test compound is connected to a training gene (yellow) vs. not (blue). (D) Model predictions of positive gene-compound pairings from test set (as log-scaled percentiles) by number of training genes connected to the test compound: performance improves with more connections.

In leave-out-compound, multiple compounds targeting a specific gene typically exert the same effect, inhibiting or activating the gene’s protein product, making them functionally related. While exceptions such as agonist-antagonist pairs exist, they are infrequent in our dataset.

In contrast, leave-out-gene presents a scenario where a compound targeting multiple genes may interact with functionally diverse targets, due to polypharmacology. The compound may hit related genes with similar protein structures, or totally unrelated genes, as different parts of its chemical structure bind to distinct targets. Our dataset was designed to mostly filter out related genes, making the second possibility more likely.

We therefore hypothesize that the asymmetry in predictive performance is attributed to the functional relatedness of targets. In leave-out-compound, compounds targeting a gene often exhibit learnably similar effects. In leave-out-gene, genes targeted by a compound are often functionally unrelated, making predicting a match for an unseen gene more difficult, as the novel gene likely has a distinct function from those in training.

To further examine the model’s behavior when gene-compound pairs are randomly left out, we employed the *leave-out-pair* strategy (Fig. 4). Here, the model accurately predicts a gene-compound pairing only if both the gene and compound have been encountered in training in a positive pair. This bias towards both the gene and the compound to have positive training set connections suggests leave-out-pair yields a less effective model than leave-out-compound.

**Figure 4.**
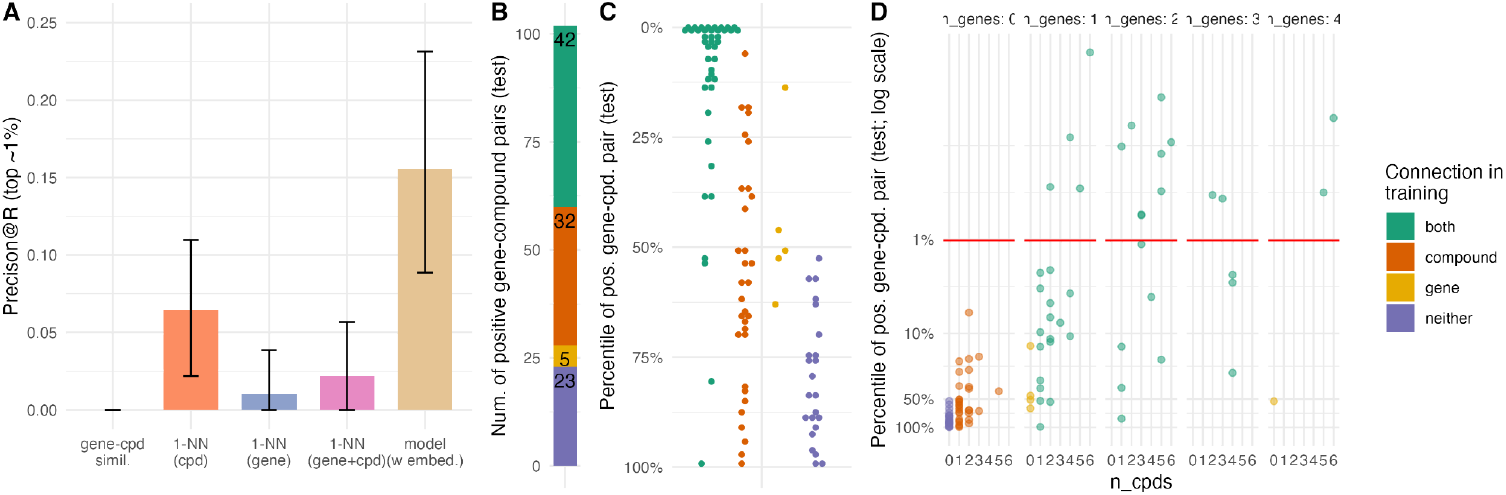
Leave-out-pair performance. (A) Proposed method (“model (w. embed.)”) vs. baselines: gene-compound similarity (“gene-cpd simil.”), 1-NN gene classifier trained for each compound (“1-NN gene”), 1-NN compound classifier trained for each gene (“1-NN compound”), and a classifier that takes a majority vote of the previous two (“1-NN gene+cpd”). Error bars: 95% CI. (B) Counts of four categories of positive gene-compound pairings from the test set: positive gene-compound pairings where the test compound is connected to a training gene (“gene”), the test gene is connected to a training compound (“compound”), both, or neither. Either the gene or the compound (or both) can be in training gene-compound pairs. (C) Model predictions of positive gene-compound pairings from test set (as percentiles): correct classification is likelier (top percentile) if both compound and gene have positive training connections (green), vs. other scenarios. (D) Model predictions of positive gene-compound pairings from test set (as log-scaled percentiles) by number of training genes connected to the test compound (panels) and training compounds connected to the test gene (x-axis): performance improves with more connections only when both – the test gene and the test compound – have positive training connections.

## Discussion

This study demonstrates the effectiveness of incorporating Cell Painting profiles of genetic perturbations to enhance drug target prediction, versus using profiles from annotated compounds only. By integrating gene morphological representations into a transformer-based model, we capture complex gene-compound relationships and improve classification performance compared to baseline methods.

Implementing this approach has the potential to accelerate drug discovery and reduce costs. Even incremental improvements in compound target prediction can have a substantial impact, especially when combined with orthogonal data sources. We estimate that creating a much larger version of the dataset we used in this study, with one million compounds and the knockout of all human genes in a single cancer cell line, would cost less than five million USD, highlighting the cost-effectiveness.

While we focused on predicting targets for novel compounds, predicting drugs for genes without known targeting compounds is equally valuable in drug discovery. However, the asymmetry in performance between leave-out-compound and leave-out-gene strategies suggests that predicting compounds that target unseen genes’ protein products is a harder learning problem, likely due to the diversity of proteins targeted by many if not most compounds. Improving the model’s ability to predict compounds that target proteins produced by unseen genes requires more diverse and extensive training data.

Future research should explore alternative embedding techniques and integrate additional data sources, such as chemical- and protein-structure based methods for gene-compound binding prediction. Further, recently curated datasets such as RxRx3-core (https://www.rxrx.ai/rxrx3-core) and MOTIVE ^12^ contain more annotated gene-compound pairs, along with corresponding Cell Painting profiles, that could be used to evaluate our method.

In conclusion, our findings underscore the value of gene embeddings in drug target prediction and highlight the need for larger, diverse datasets to handle novel genes. Advancing these methods and integrating complementary data sources can accelerate the identification of potential drug targets and facilitate the development of new therapeutic interventions, streamlining the drug discovery process.

## Methods

### Data collection and preprocessing

The CPJUMP1 dataset ^8^ was used in this study, which contains Cell Painting profiles of chemical and genetic perturbations. The dataset was designed to include pairs of compounds and genes expected to have matching relationships. Chemical perturbations were selected from the Drug Repurposing Hub dataset ^10^, while genetic perturbations involved either gene overexpression using open reading frames (ORFs) or gene inhibition via CRISPR-Cas9-mediated knockout.

Compound selection criteria included targeting genes from diverse families, having at least two compounds targeting each gene, and excluding non-selective or less potent compounds. For each gene, one ORF, two CRISPR guides, and one or two known compounds influencing the gene product’s function were chosen. The dataset contains 302 compounds and 160 genes, fitting into three 384-well plates with controls. Positive and negative controls for each perturbation modality were included. We note that the original paper ^8^ mentions 303 compounds; however, depending on how the SMILES are standardized, we end up with 302±1 unique compounds. In this paper, we consider the dataset to have 302 unique compounds.

Perturbation profiles were generated by either transfecting cells with the respective genetic reagents (for CRISPR and ORF) or by treating with compounds, and then using the Cell Painting protocol ^13^ to fix, stain, and image the cells.

The resulting images were processed using the CellProfiler software to extract morphological features at the single-cell level. The subsequent data processing steps were performed using PyCytominer ^14^. First, the single-cell profiles were aggregated by computing the median profile for each well. The aggregated profiles were then normalized by subtracting the median and dividing by the median absolute deviation (m.a.d.) of each feature, using the median and m.a.d. of the negative control wells on the plate. Finally, redundant features (such that no pair of features has Pearson correlation greater than 0.9) and features with near-zero variance across all plates were filtered out. Feature selection was performed on each plate separately, and then the intersection of features across all plates was used as the final list of features (n=100 features). An alternative approach would have been to perform feature selection across all plates simultaneously; however, early experiments indicated that, given the high redundancy of features, the chosen approach yielded satisfactory results. The complete data processing workflow for producing well-level profiles is available at https://github.com/jump-cellpainting/pilot-cpjump1-data/tree/v1.0.0. The embedding of a perturbation is obtained by computing the median across all the replicates of the perturbation.

For gene embeddings, we considered using only the ORF profile, only the CRISPR profiles (with maximum or average predictions across guides), or both ORF and CRISPR profiles (again with maximum or average predictions across CRISPR guides). Our previous work ^8^ showed that CRISPR and ORF profiles for the same gene are often uncorrelated, rather than anti-correlated as might be expected. This lack of correlation raises questions about the appropriateness of combining these data types. However, we hypothesize that the uncorrelated profiles may capture complementary aspects of gene function. Our transformer model is designed to extract relevant information from each perturbation type without requiring them to be correlated or anti-correlated. While the biological basis for this lack of correlation remains unclear, using both CRISPR and ORF profiles may provide a more comprehensive representation of gene function. Based on this rationale, we chose to use both ORF and CRISPR profiles, and averaged predictions across CRISPR guides.

The full CPJUMP1 dataset includes ORF, CRISPR, and compound profiles for two human cancer cell lines (U2OS and A549) at two different timepoints post-treatment. For the experiments presented in this paper, we used a subset of the data, focusing on the U2OS cell line and the timepoints that were, at the time, considered the most likely to be used in the full experiment by the JUMP Cell Painting Consortium. Specifically, we used data from U2OS cells treated with compounds for 24 hours, CRISPR-perturbed for 144 hours, and ORF-perturbed for 48 hours.

### Model development

We formulated the prediction of a compound’s gene target as a translation problem and employed a transformer model to solve it. Transformer models, originally introduced for natural language processing tasks, are neural network architectures that utilize self-attention mechanisms. These models are particularly suitable for processing data with complex dependencies, as they model ^15^ the relationships among all input data elements simultaneously rather than in a specific order.

The transformer model consists of an encoder and a decoder. The gene embeddings (CRISPR, ORF, or CRISPR+ORF) are fed into the encoder, while the compound information is input to the decoder (Fig 1c). The encoder processes the gene embedding and generates a fixed-length vector representation, called the context vector, which captures the essential features of the gene embedding required for the translation task.

The decoder then uses the context vector and the previously generated target data (compound profiles) to predict the gene target of a compound. By leveraging both the information in the context vector and the compound profiles, the decoder produces the final prediction, establishing a mapping from gene targets to compounds.

The transformer model’s output is passed through a fully connected layer and a softmax layer to obtain a binary prediction probability indicating the likelihood of the gene being a target of the compound. This architecture learns a similarity metric between pairs of samples, optimizing the similarity among similar samples and minimizing it among dissimilar ones.

### Model training and evaluation

The dataset was split into training (60%), validation (20%), and test (20%) sets using a stratified sampling approach to ensure a balanced representation of gene-compound pairs across the splits. The training set was used to train the model, the validation set for hyperparameter optimization, and the test set for final performance evaluation. The PyTorch machine learning library was used for model implementation, with the dataset converted into DataLoader objects with a batch size of 1024. Data within each set was randomly shuffled and grouped into batches.

Hyperparameter tuning was performed using a grid search over the validation set to identify the best combination of parameters for the transformer model. The search space included batch size, number of attention heads, dropout rate, learning rate, and the number of training epochs. Our framework employed three complementary metrics: cross-entropy loss for training optimization, precision-recall AUC on the validation set for early stopping to prevent overfitting, and Precision@R for final model selection on the validation set. Precision@R, defined as the proportion of true positives among the top R ranked predictions (where R is the total number of true positives in the dataset), measures the model’s ability to identify promising compound candidates.

The final model was then trained using these optimized hyperparameters and evaluated on the test set to assess its generalization ability using the Precision@R metric.

| Parameters | Search Space |
| --- | --- |
| Number of attention heads (nhead) | 4, 5, 10 |
| Number of hidden (nhid) | 128, 512, 1024 |
| Number of layers (nlayers) | 4, 6 |
| Dropout rate | 0.1, 0.2 |
| Learning rate | 0.001, 0.0001 |

## Acknowledgments

The authors gratefully acknowledge a grant from the Massachusetts Life Sciences Center Bits to Bytes Capital Call program for funding the data production. We appreciate funding to support data analysis and interpretation from members of the JUMP Cell Painting Consortium.

Researchers in the Carpenter–Singh lab were supported in part by NIH R35 GM122547 to AEC. The authors appreciate helpful comments from Erin Weisbart, Esteban Miglietta, and members of the JUMP Cell Painting Consortium.

## Author contributions

N.S.I., D.J.M., S-Y G.C, J.A., and S.S. wrote the code and conducted the analysis; N.C. provided guidance on the CPJUMP1 dataset, S.S., P.R., and A.E.C. supervised the research. All authors designed the experiments and wrote and edited the paper.

## Data and code availability

All code to reproduce this analysis is available at

https://github.com/carpenter-singh-lab/2024_Iyer_Genemod. All the corresponding data is available as part of the *cpg0000-jump-pilot* dataset ^16^, available from the Cell Painting Gallery ^17^ on the Registry of Open Data on AWS (https://registry.opendata.aws/cellpainting-gallery/).

## Declaration of interests

The Authors declare the following competing interests: S.S. and A.E.C. serve as scientific advisors for companies that use image-based profiling and Cell Painting (A.E.C: Recursion, SyzOnc, Quiver Bioscience, S.S.: Waypoint Bio, Dewpoint Therapeutics, Deepcell) and receive honoraria for occasional talks at pharmaceutical and biotechnology companies. All other authors declare no competing interests.

## Notes

### Summary of Updates

- Clarified data preprocessing, model training, and evaluation methodology in the Methods section - Added new Supporting Information for over-representation analysis of gene-compound predictions, details on the molecules and targets studied, analysis of chemical diversity among compounds targeting individual genes, and model convergence analysis - Made code repository public for reproducibility and provided details on hyperparameter tuning

