## Supporting Information for "Cell morphological representations of genes enhance prediction of drug targets"

#### A. Over-Representation Analysis of Gene-Compound Predictions

The top ~1% (n=89) of gene-compound pairs, ranked by probability of match, were examined for overrepresented genes and pathways. Gene enrichment analysis (Table A1) was performed on the 18 genes with at least two known compound connections, comparing their occurrence in correctly predicted vs. incorrectly predicted connections. Pathway enrichment analysis (Table A2) was conducted by mapping genes to pathways using KEGG[1] and WikiPathways[2] annotations. Neither showed significant enrichment after multiple testing corrections, suggesting the model's predictions were not biased towards specific genes or pathways.

| **Gene** | **Correctly predicted** | **Set Size** | **P-value** | **FDR** |
|---|---:|---:|---:|---:|
| TUBB | 2 | 2 | 0.016 | 0.142 |
| IGF1R | 2 | 2 | 0.016 | 0.142 |

*Table A1: Gene enrichment analysis results for the top 89 gene-compound pairs. Results shown are filtered by a false discovery rate (FDR) < 0.99. No significant gene enrichment was observed after multiple testing correction.*

| **Pathway** | **Source** | **Correctly predicted** | **Set Size** | **P-value** | **FDR** |
|---|---|---:|---:|---:|---:|
| Pathogenic Escherichia coli infection | Wiki | 3 | 3 | 0.007 | 0.625 |
| Parkin-Ubiquitin Proteasomal System pathway | Wiki | 3 | 3 | 0.007 | 0.625 |

*Table A2: Pathway enrichment analysis results for the top 89 gene-compound pairs. Results shown are filtered by a false discovery rate (FDR) < 0.99. No significant pathway enrichment was observed after multiple testing correction.*

### B. Details of genes and compounds profiled in the CPJUMP1 dataset.

The CPJUMP1 dataset, introduced by Chandrasekaran et al. [3], comprises Cell Painting profiles from chemical and genetic perturbations in U2OS (osteosarcoma) and A549 (lung carcinoma) cell lines. The dataset includes 302 small molecules from the Drug Repurposing Hub [4] and covers 160 genes through overexpression via open reading frames (ORFs) and inhibition using CRISPR-Cas9-mediated knockout. In brief, these gene-compound pairings were selected based on criteria such as targeting proteins encoded by diverse gene families, having at least two compounds targeting each gene product, and excluding non-selective or insufficiently potent compounds. The dataset also includes negative and positive controls for each perturbation modality.

Comprehensive details on the genes and compounds profiled, as well as the experimental conditions used, can be found in the original paper [3] and its accompanying GitHub repository (https://github.com/jump-cellpainting/2024_Chandrasekaran_NatureMethods). The repository at https://github.com/jump-cellpainting/JUMP-Target/tree/v1.1.1 contains metadata related to the chemical and genetic perturbations tested and the plate layouts used.
All compound annotations were obtained from the Broad Repurposing Hub (https://repo-hub.broadinstitute.org/repurposing#download-data). Specifically, the annotations were sourced from the file https://s3.amazonaws.com/data.clue.io/repurposing/downloads/repurposing_drugs_20200324.txt. It is important to note that several compounds are annotated with multiple targets, and the full list of targets for each compound is provided in the "target_list" column of the compound metadata https://github.com/jump-cellpainting/JUMP-Target/blob/v1.1.1/JUMP-Target-1_compound_metadata.tsv.

#### C. Analysis of chemical diversity among compounds targeting individual genes.

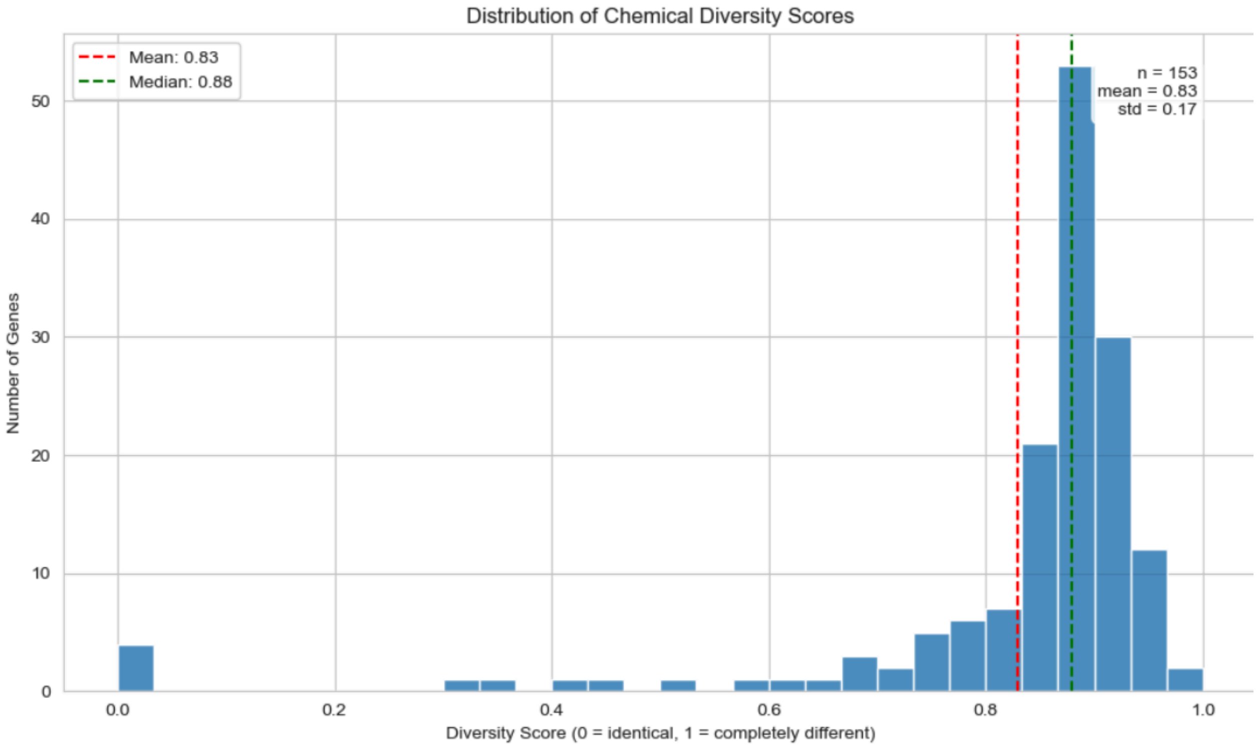


*Figure C1: For each gene, we calculated a diversity score (ranging from 0 to 1) based on pairwise Tanimoto similarities of Morgan fingerprints among its targeting compounds, where 0 indicates identical compounds and 1 indicates completely different structures. The distribution across all genes (n=153; genes that have at least two compounds targeting them, and for which we have ORF/CRISPR Cell Painting profiles) shows a mean diversity score of 0.83 (SD=0.17), indicating that most genes are targeted by structurally diverse compounds. The right-skewed distribution with most scores between 0.8-0.9 suggests our compound collection generally maintains good chemical diversity within gene-specific compound sets.*

### D. Model Training and Convergence Analysis

To validate model fitting behavior, we analyzed the convergence characteristics across all training runs (n=72) of the leave-out-compound formulation. Individual learning curves (Figure D1) reveal distinct behavioral patterns: some runs exhibit classic divergence between training and validation loss, while others show minimal learning progress (flat loss curves).

Note that our framework employed three complementary metrics: cross-entropy loss for training optimization, precision-recall AUC on the validation set for early stopping to prevent overfitting (Figure D2), and Precision@R (Figure D3) for final model selection on the validation set.

The configuration we selected as optimal is highlighted in yellow. While the learning curves might suggest suboptimal performance, this interpretation requires important context: The initial loss value (step 0, not shown in figures) is 0.73 ± 0.11 (SD) and 0.72 ± 0.09 (SD), for training and validation, respectively, across all runs. Significant learning occurs in the earliest training steps, with loss values dropping to 0.079 ± 0.012 (SD) and 0.046 ± 0.002 (SD), for training and validation, respectively, by step 1. This rapid initial improvement suggests effective model training despite the apparent plateau in later steps.

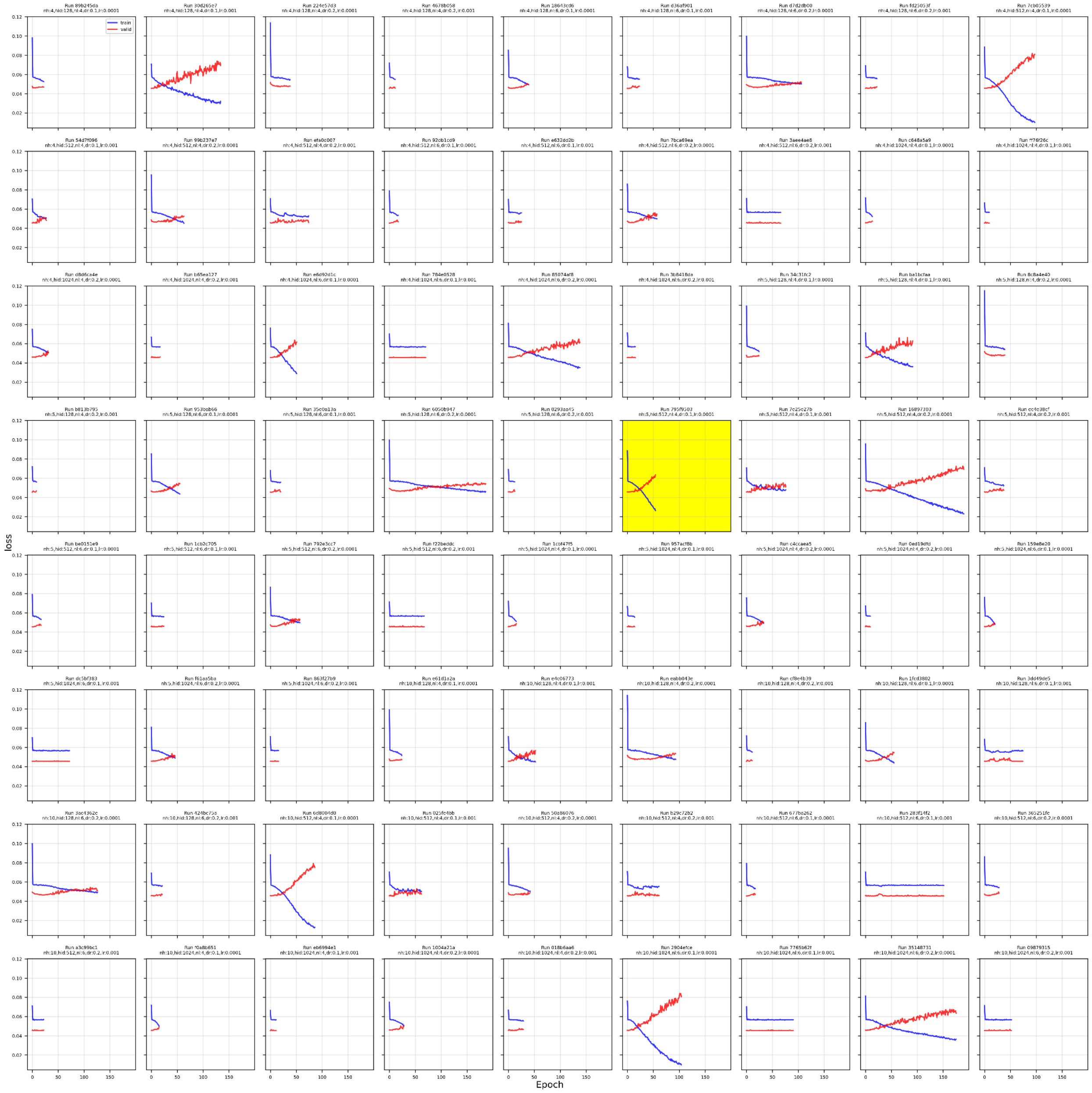


*Figure D1: Individual learning curves (cross-entropy loss) for all 72 training runs showing training (blue) and validation (red) loss across epochs.*

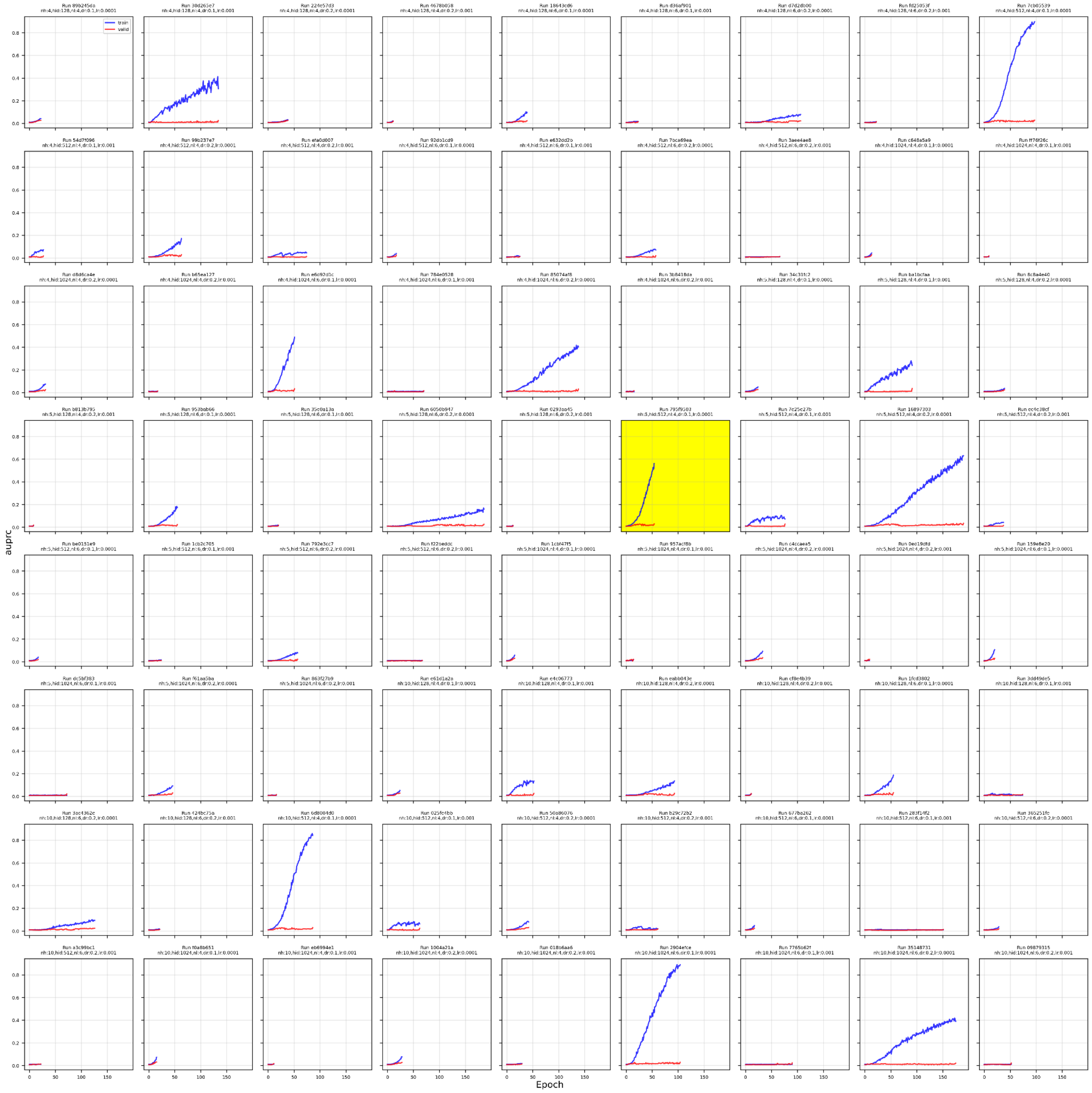


*Figure D2: Individual learning curves (PR-AUC) for all 72 training runs showing training (blue) and validation (red) loss across epochs.*

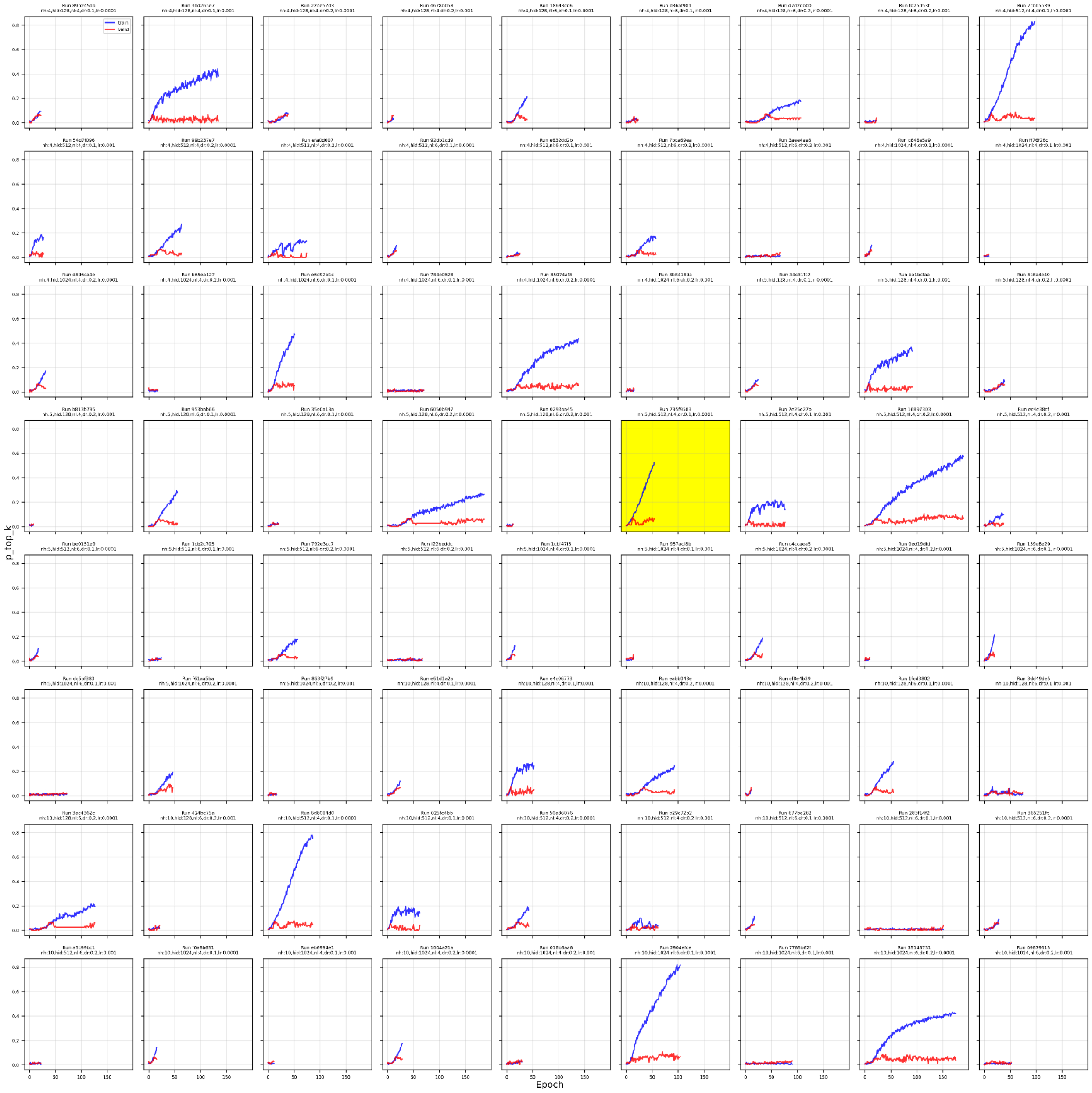

*Figure D3: Individual learning curves (Precision@R) for all 72 training runs showing training (blue) and validation (red) loss across epochs.*
